## Supplementary Figures for "Screening non-conventional yeasts for organic acid tolerance and engineering *Pichia occidentalis* for production of *cis*,*cis*-muconic acid"

### Supplementary Results

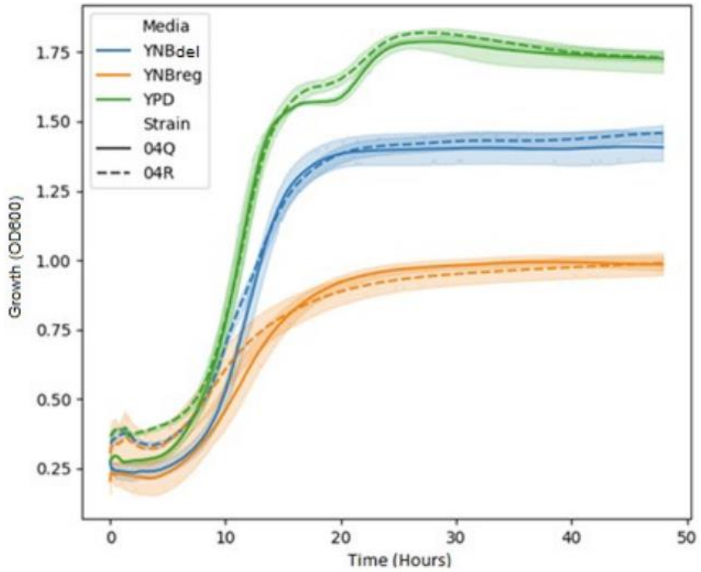


**Supplementary Figure 1. Comparison of *P. occidentalis* strain 04Q (YB-3389) and 04R (Y-6545) growth in standard 1× YNB medium (YNBreg) vs growth in 3× YNB medium (YNBdel).** Highlighted area shows the standard deviation of $n$ = 3 independent biological samples. YNBreg contains 20 g $L^{-1}$ glucose, 5.1 g $L^{-1}$ ammonium sulfate and 1.7 g $L^{-1}$ Yeast Nitrogen Base. YNBdel contains 20 g $L^{-1}$ glucose, 5.1 g $L^{-1}$ ammonium sulfate and 5.1 g $L^{-1}$ Yeast Nitrogen Base.

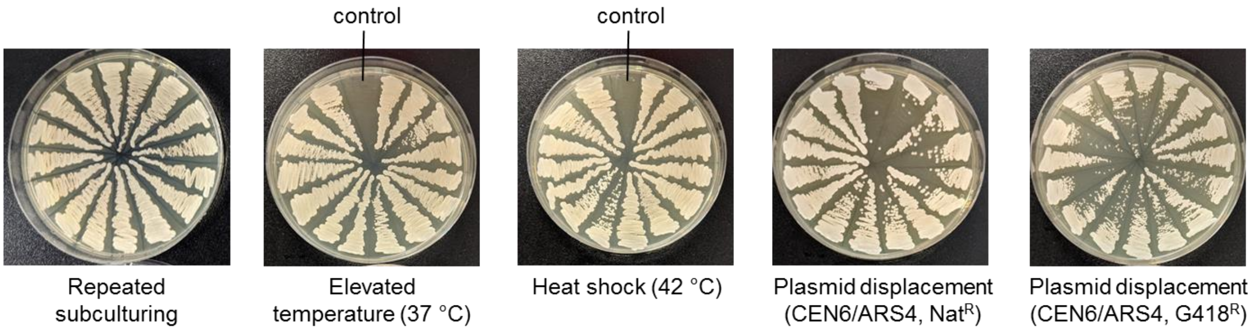


**Supplementary Figure 2. Attempted curing of plasmid pCas-Hyg-CEN6ARS4-ADE2 from *P. occidentalis* Y-7552.** Various curing methods were assessed to cure cells of the pCas-Hyg-ADE2 plasmid. Hyg$^R$ colonies were subcultured six times without selection at 30 °C (repeated subculturing) or 37 °C (elevated temperature) prior to screening colonies on YPD agar plates containing hygromycin. Curing was also attempted by heat-shocking cells at 42 °C in a mock lithium-acetate-PEG transformation (heat shock) or by attempting to displace the original pCas-Hyg-ADE2 plasmid with Nat$^R$ or G418$^R$ derivatives containing the same CEN6/ARS4 origin. Wild-type *P. occidentalis* lacking a pCas plasmid was included as a control.

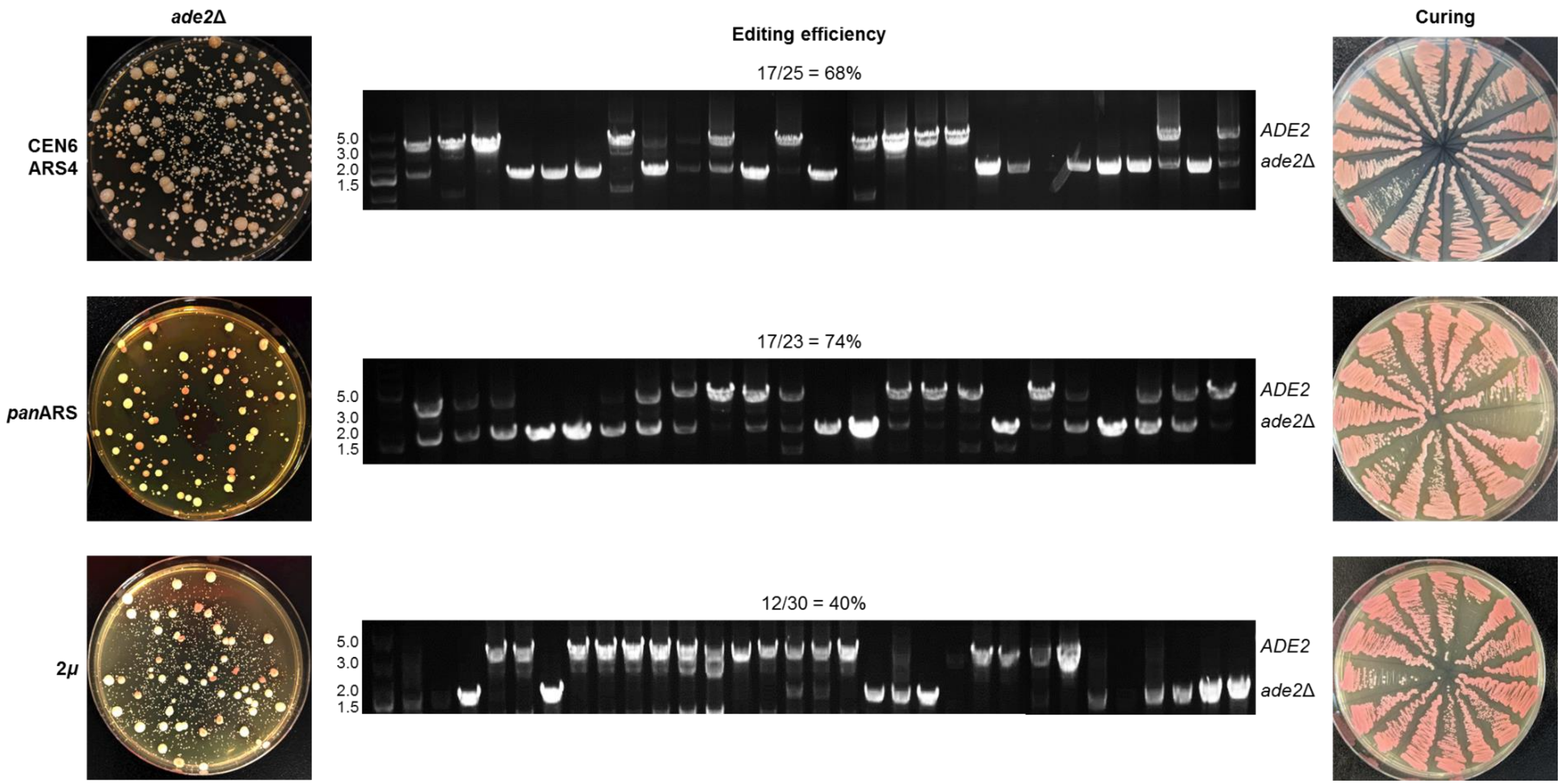


**Supplementary Figure 3. Deletion of *ADE2* in *P. occidentalis* Y-7552 using various pCas plasmid origins.** The *P. occidentalis ADE2* gene was deleted using a pCas-Hyg-ADE2 plasmid containing a CEN6/ARS4, *pan*ARS, or 2*μ* origin. Editing efficiency is shown based on screening of 23-30 random colonies. One *ade2*Δ mutant colony containing each plasmid origin was subcultured six times in YPD without selection and resultant colonies were screened for loss of Hyg$^R$ on YPD agar plates containing hygromycin. Repeating *ADE2* deletion using different plasmid origins routinely yielded similar editing efficiencies.

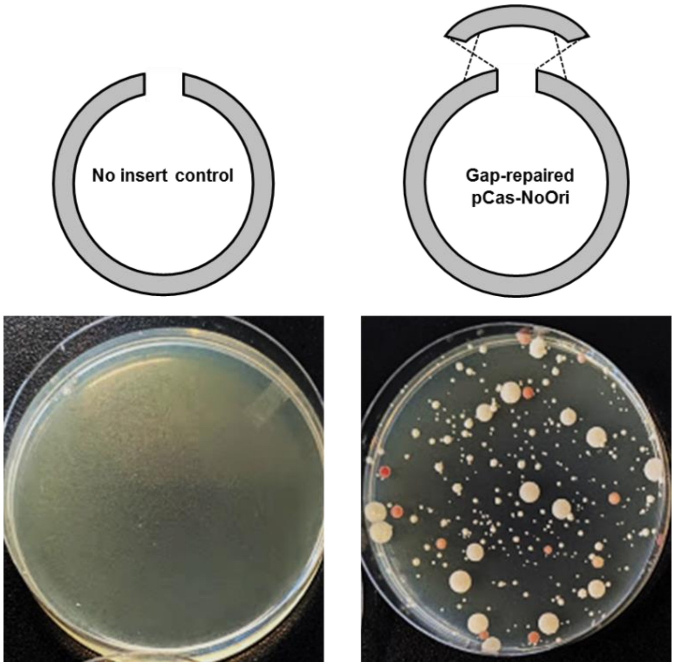


**Supplementary Figure 4. Deletion of *ADE2* in *P. occidentalis* Y-7552 without a plasmid origin.** The *P. occidentalis ADE2* gene was deleted using a pCas-Hyg-ADE2 plasmid lacking a plasmid origin. Plasmid pCas-Hyg-ADE2 was digested within the CEN6/ARS4 origin and the linearized plasmid was used to transform *P. occidentalis* along with an overlapping gap repair template lacking a plasmid origin. The resulting origin-less pCas plasmid generated red colonies following selection on YPD agar containing hygromycin. Omitting a linear gap repair template failed to generate $Hyg^R$ transformants (no insert control).

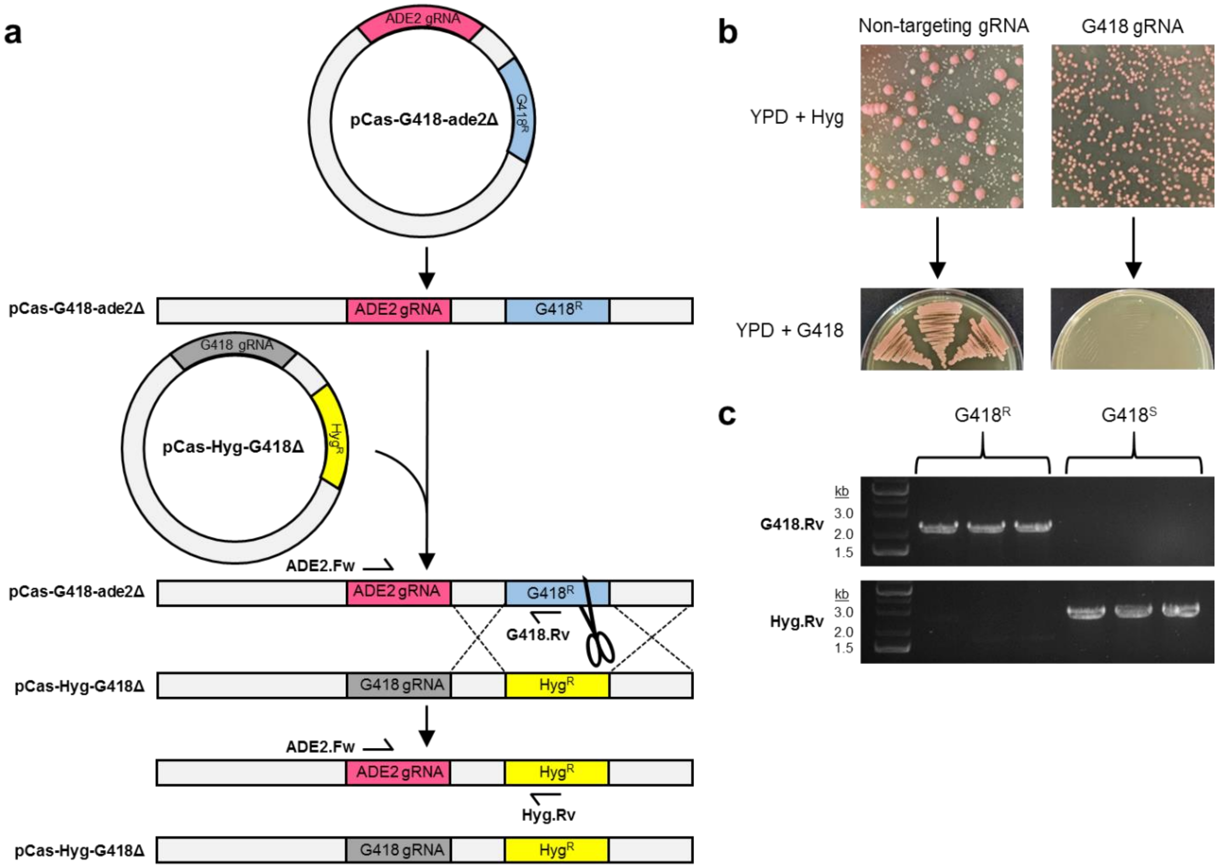


**Supplementary Figure 5. Plasmid integration and antibiotic marker recycling in *P. occidentalis* Y-7552. a,** Proposed mechanism of plasmid integration and antibiotic marker recycling. Introduction of plasmid pCas-G418-ade2Δ and selection of G418$^R$ colonies yields pink *ade2*Δ colonies upon integration of pCas-G418-ade2Δ. Transformation of a pink G418$^R$ *ade2*Δ mutant with plasmid pCas-Hyg-G418Δ harboring a Hyg$^R$ marker and a gRNA targeting chromosomal G418$^R$ yields Hyg$^R$ colonies upon chromosomal integration of pCas-Hyg-G418Δ. Transcription of the G418 gRNA introduces a double stranded DNA break to the chromosomal G418$^R$ marker, which is repaired by the homologous chromosomal Hyg$^R$ marker. The resulting strain lacks a G418$^R$ marker. Primers used for PCR screening in **c** are shown. **b,** Introduction of a gRNA targeting a chromosomal G418$^R$ marker yields a substantial increase in Hyg$^R$ transformants compared to a control transformation utilizing a non-targeting gRNA (top). In line with the mechanism outlined in **a**, Hyg$^R$ colonies transformed with G418 gRNA lost G418$^R$, while colonies transformed with a non-targeting gRNA retained G418$^R$ (bottom). **c,** Colony PCR confirmation of the proposed marker swapping mechanism outlined in **a**. Screening G418$^S$ colonies using primers ADE2.Fw + G418.Rv confirms loss of the chromosomal G418-resistance marker. Screening G418$^S$ colonies using primers ADE2.Fw + Hyg.Rv confirms acquisition of the Hyg$^R$ marker not observed in G418$^R$ colonies transformed with a non-targeting gRNA. Three representative G418$^R$ and G418$^S$ colonies were screened.

*fcy1*Δ *fcy1*Δ

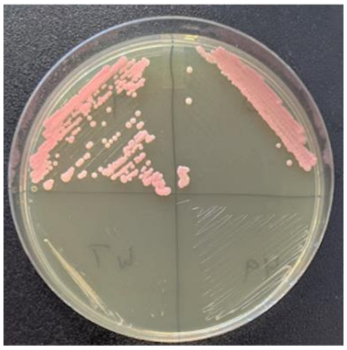

WT WT

**Supplementary Figure 6. Deletion of *FCY1* confers resistance to 5-fluorocytosine (5-FC).** *FCY1* was deleted in an *ade2*Δ G418$^{R}$ host using pCas-Hyg-CEN6ARS4-PoADE2. Transformants were restreaked onto YPD agar plates overlaid with 400 µl of a 10 g $L^{-1}$ solution of 5-FC.

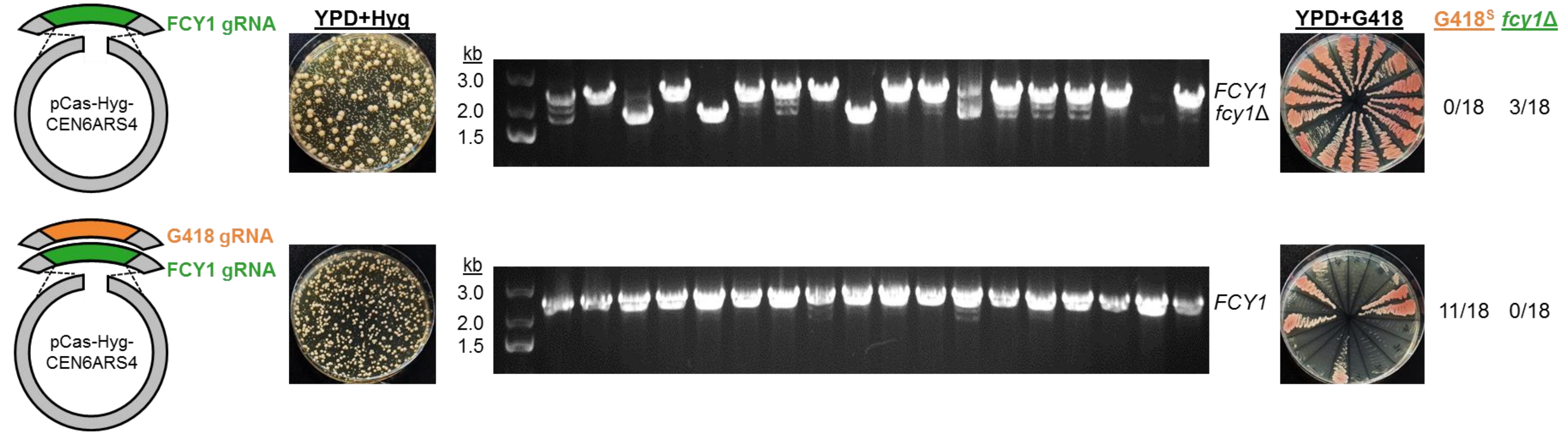


**Supplementary Figure 7. Deletion of *FCY1* and G418$^{R}$ marker recycling in *P. occidentalis* Y-7552 using dual gRNA species.** Dual gRNA species targeting a chromosomal G418$^{R}$ marker and the *FCY1* gene were introduced to a G418$^{R}$ *ade2*Δ mutant (bottom) and compared to a control transformation using a single *FCY1* gRNA (top). Hyg$^{R}$ transformants were scored for *fcy1*Δ by colony PCR and G418$^{S}$ by restreaking transformants onto YPD+G418.

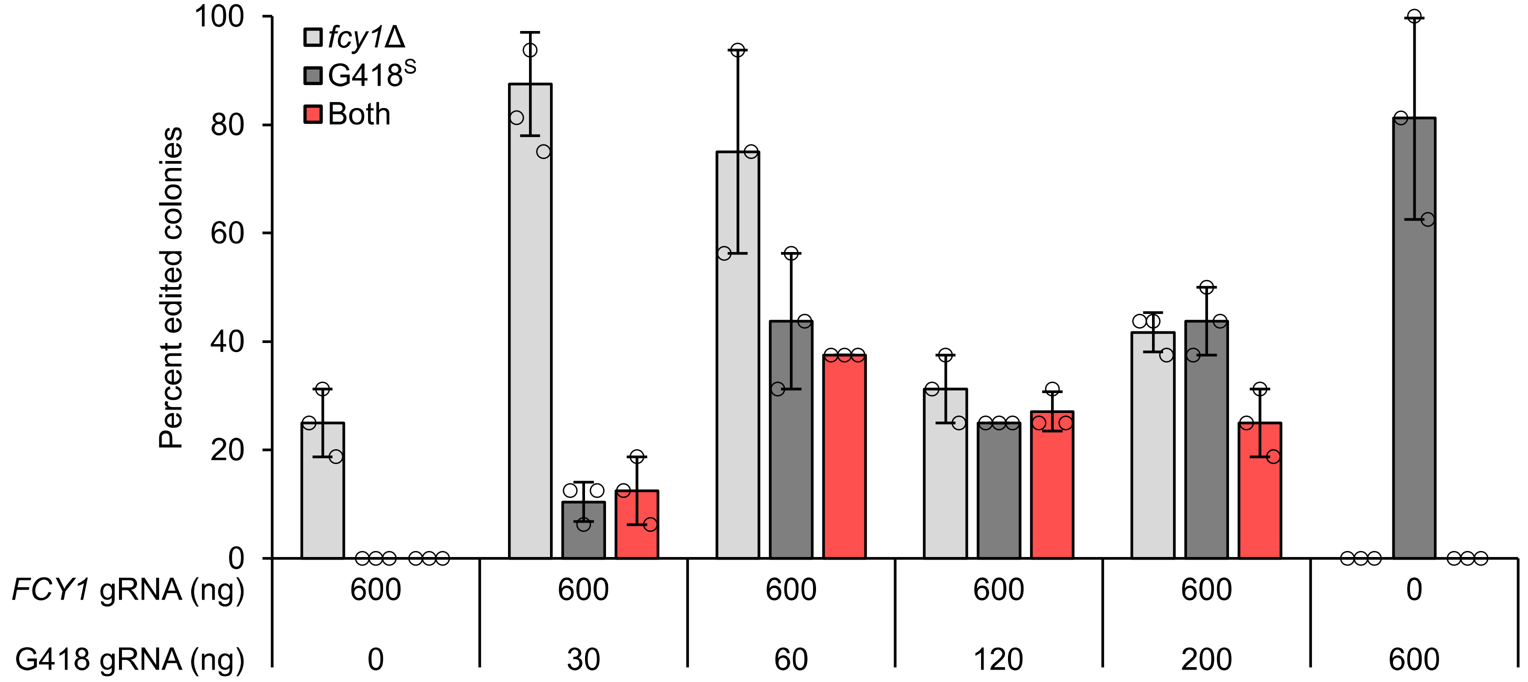


**Supplementary Figure 8. Titration of *FCY1* and G418$^{R}$ gRNA species.** Varying amounts of G418 gRNA and 600 ng of *FCY1* gRNA were introduced to a G418$^{R}$ *ade2*Δ mutant using pCas-Hyg-CEN6ARS4. Sixteen Hyg$^{R}$ transformants from each condition were scored for *fcy1*Δ and G418$^{S}$ by restreaking onto YPD plates containing 5-FC and G418, respectively. Error bars represent the mean ± s.d. of $n$ = 3 independent biological samples. Source data underlying Fig. S8 are provided in a Source Data file.

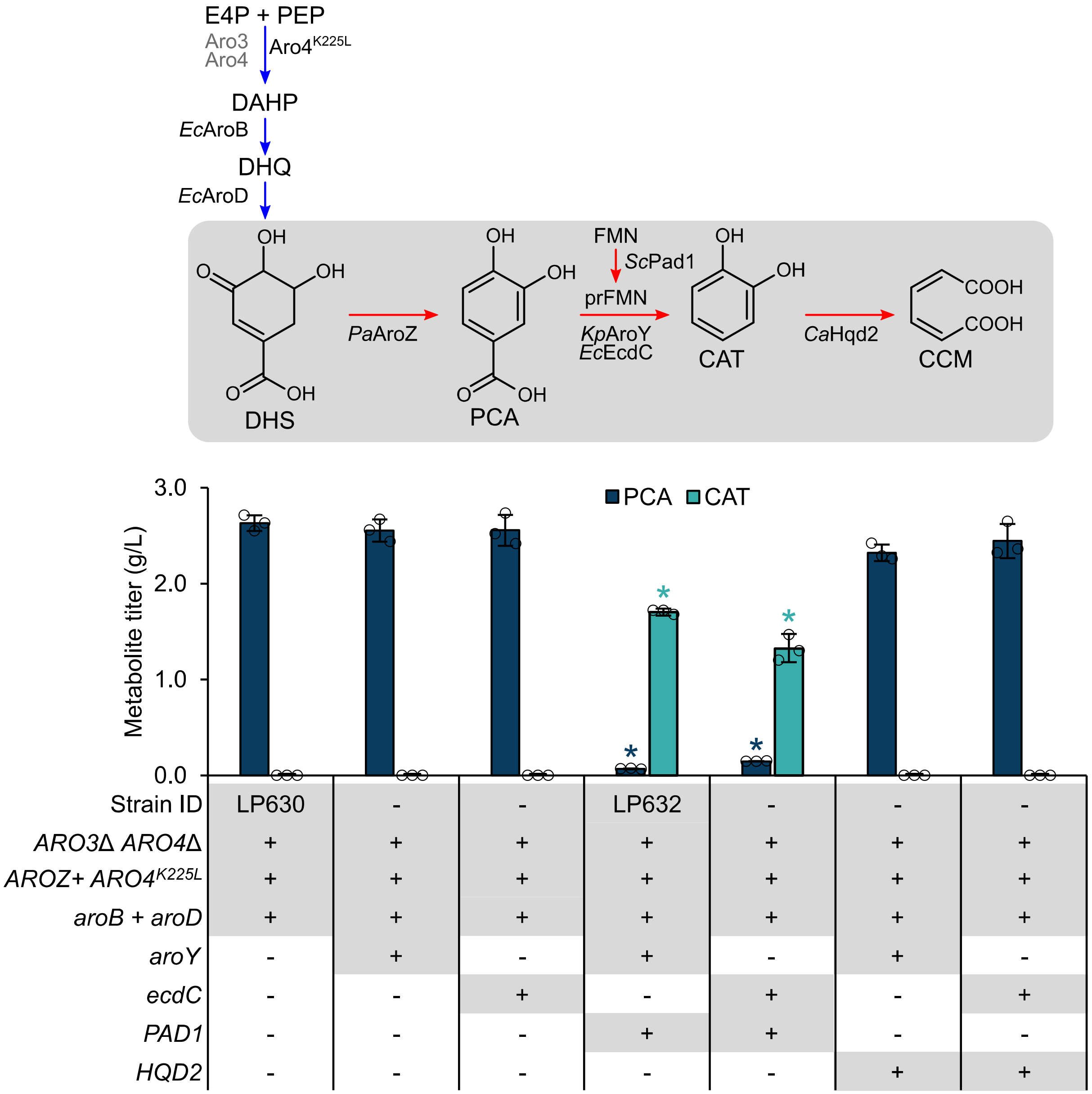


**Supplementary Figure 9. Pad1 is required for PCA decarboxylase activity in engineered *P. occidentalis*.** Combinations of PCA decarboxylase (*Kp*AroY or *Ec*EcdC), FMN prenyltransferase (*Sc*Pad1) and catechol dioxygenase (*Ca*Hqd2) were introduced to PCA-producing *P. occidentalis* (LP630). The native shikimate and heterologous CCM synthesis pathway are depicted in blue and red, respectively. Asterisks (*) denote a significant decrease in PCA titer or increase in catechol titer relative to the parent strain ($P < 0.05$). Statistical differences between parent and derivative strains were tested using two-tailed Student's *t*-test. Error bars represent the mean ± s.d. of $n = 3$ independent biological samples. Abbreviations: CAT, catechol; CCM, *cis*,*cis*-muconic acid; DAHP, 3-deoxy-D-arabinoheptulosonate 7-phosphate; DHQ, 3-dehydroquinate; DHS, 3-dehydroshikimate; E4P, erythrose 4-phosphate; FMN, flavin mononucleotide; PCA, protocatechuic acid; PEP, phosphoenolpyruvate; prFMN; prenylated flavin mononucleotide. Source data underlying Fig. S9 are provided in a Source Data file.

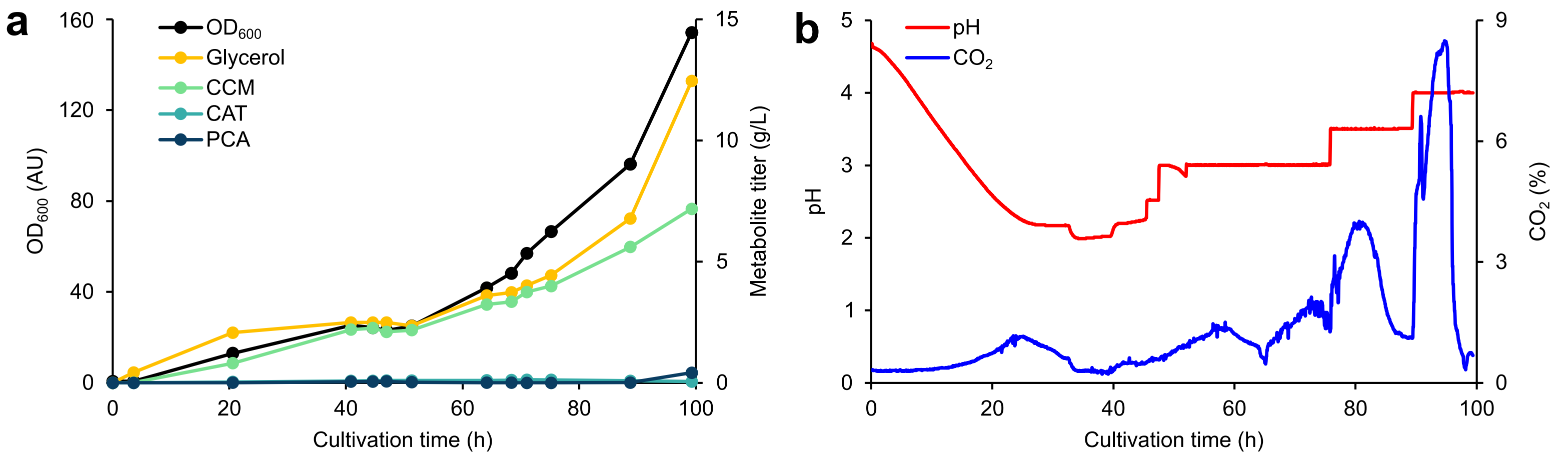


**Supplementary Figure 10. Cultivation of a CCM-producing *P. occidentalis* strain (LP635) in a glucose-limited fed-batch fermentor at low pH.** A CCM-producing strain (LP635) was fed 0.5 L of medium containing 360 g $L^{-1}$ glucose. Biomass ($OD_{600}$) and heterologous product accumulation (**a**), and pH and $CO_2$ traces (**b**) are depicted. pH was adjusted manually using 4 N NaOH based on a decline in $CO_2$ production. Source data underlying Fig. S10 are provided in a Source Data file.

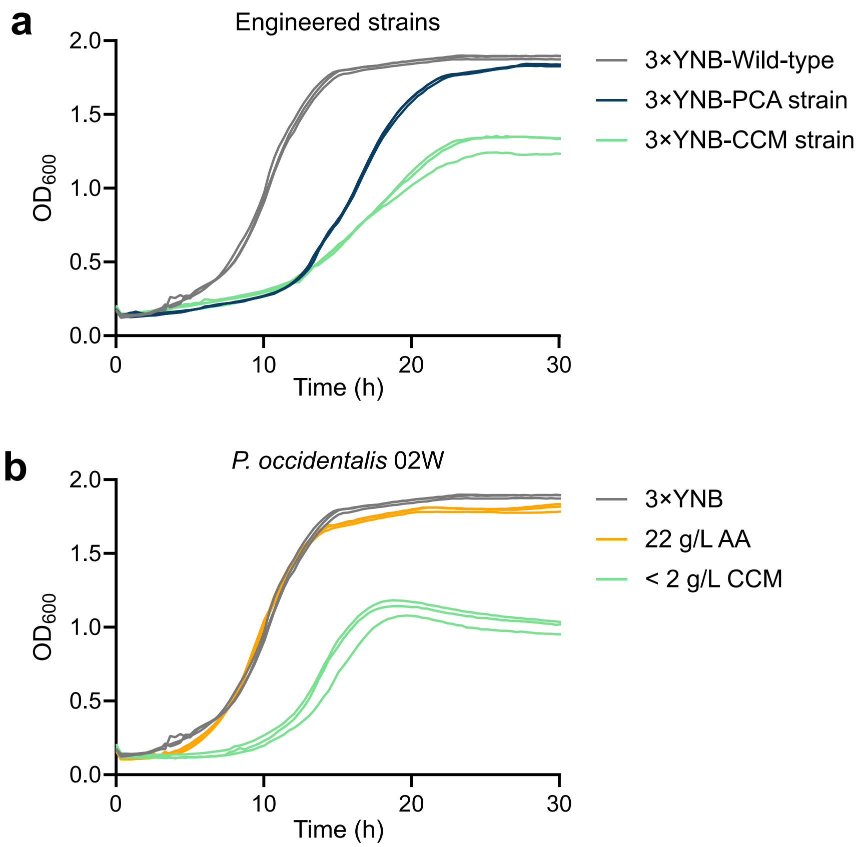


**Supplementary Figure 11. Growth curves of engineered CCM-producing *P. occidentalis* and the wild-type strain supplemented with exogenous CCM. a,** Growth of CCM-producing *P. occidentalis* and its PCA-producing precursor. Strains were cultivated in 3× YNB. **b,** Growth of wild-type *P. occidentalis* in 3× YNB saturated with adipic acid (22 g $L^{-1}$ or 0.15 M) or CCM (< 2 g $L^{-1}$). $n$ = 3 independent biological samples are overlaid for each strain and growth condition. Source data underlying Fig. S11 are provided in a Source Data file.

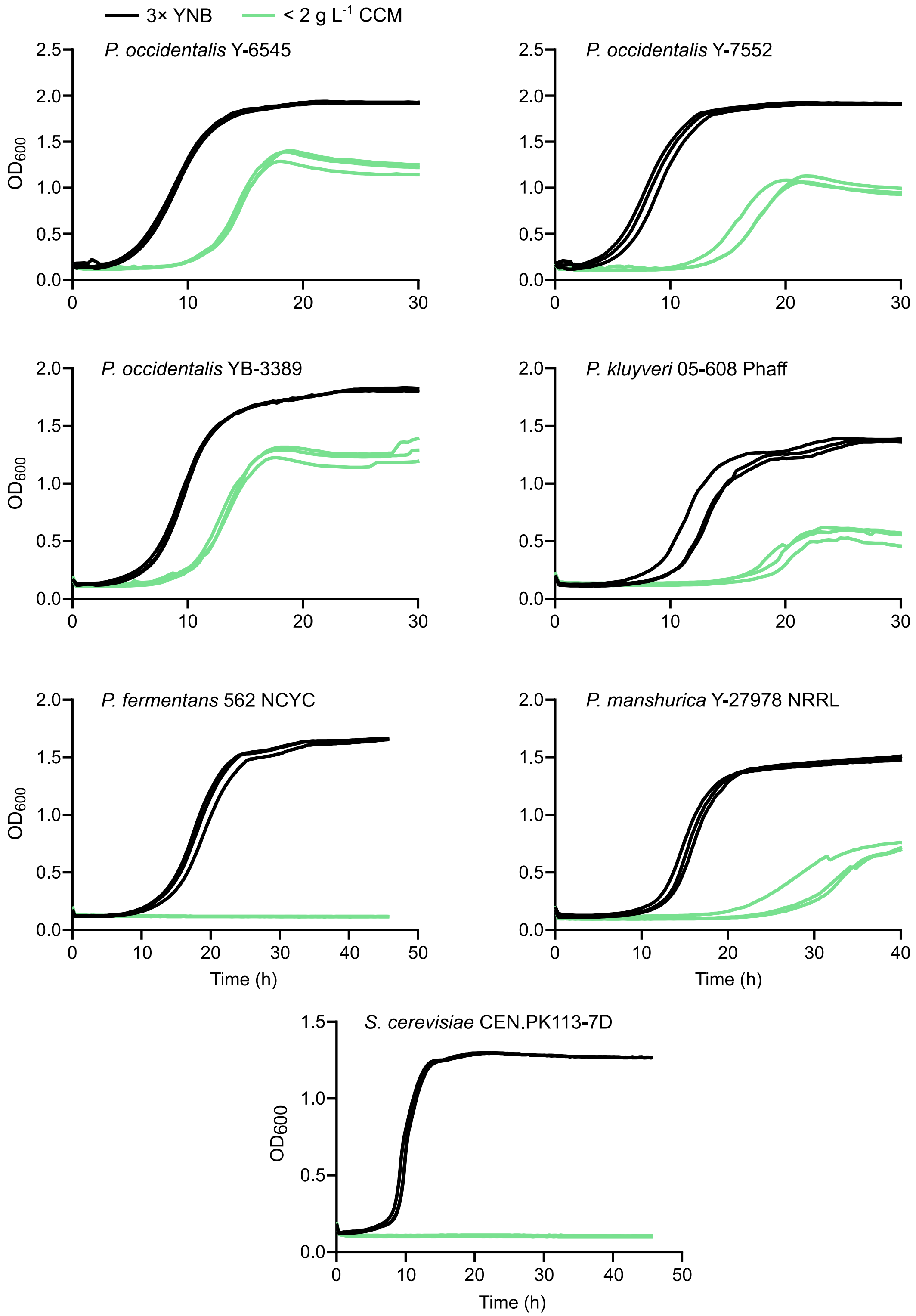


**Supplementary Figure 12. Growth curves of adipic-acid-tolerant *Pichia* strains supplemented with exogenous CCM.** 3× YNB was saturated with CCM (< 2 g $L^{-1}$). $n$ = 3 independent biological samples are overlaid for each strain and growth condition. Source data underlying Fig. S12 are provided in a Source Data file.
